# RAGulate: Retrieval-Augmented Generation for Post-hoc Literature-Grounded Regulatory Assessment

**DOI:** 10.64898/2026.01.20.700704

**Authors:** Mehrdad Zandigohar, Yang Dai

## Abstract

**Motivation:** Prioritization of transcription factor (TF)–target relationships predicted by computational models for experimental validation often requires biologists to manually inspect heterogeneous and context-dependent evidence scattered across the biomedical literature. Large Language Models (LLMs) offer a promising solution to streamline this task. However, their reliance on general-purpose knowledge may lead to hallucinations and inaccurate interpretations.

**Results:** We present RAGulate, a retrieval-augmented generation (RAG) framework for literature-grounded assessment of transcriptional regulation. RAGulate leverages CollecTRI, an external regulatory knowledge base, and integrates alias-aware query expansion, sparse and dense retrieval, maximum-marginal-relevance re-ranking, and LLM-based classification of predictions within a modular pipeline. Using a balanced TF–target–context benchmark from the same resource, we evaluate retrieval, classification, and evidence faithfulness. While CollecTRI provides TF-target links with supporting PubMed Identifiers (PMIDs), RAGulate infers the context of each interaction from the retrieved literature. Results show that alias normalization markedly improves retrieval recall, while hybrid retrieval, which merges lexical and embedding-based candidates, achieves the highest evidence recovery across all cut-offs. Conditioning LLMs on retrieved documents consistently improves AUROC and AUPR for classifying whether a TF–target interaction is supported in the specified context compared with direct prompting. RAGulate reduces hallucinations and improves PMID-level citation correctness, producing explanations that faithfully reflect the supporting literature. RAGulate represents a knowledge-based AI tool that partners with biologists to accelerate the process of TF-target prioritization for experimental validation and foster hypothesis generation.

**Availability and implementation:** The software and tutorials are available at github.com/YDaiLab/RAGulate.

## 1 Introduction

The reconstruction of gene regulatory networks (GRNs) from multiomics data is fundamental for elucidating the mechanisms by which transcription factors (TFs) orchestrate the regulation of target genes, thereby providing critical insights into cellular function and behavior. Recent computational models can infer GRNs from single-cell RNA-seq and ATCA-seq data, producing numerous predicted TF-gene regulatory relationships. Representative tools include regulon-centric approaches such as SCENIC (Aibar *et al*. 2017), probabilistic frameworks such as BITFAM (Gao *et al*. 2021), and more recent methods including CellOracle (Kamimoto *et al*. 2023) and scRegulate (Zandigohar *et al*. 2025) that infer context-specific regulatory programs.

However, the existing GRN inference frameworks primarily focus on optimizing inference accuracy through benchmarking on simulated datasets. When applying to real-world data, these methods typically generate tens of thousands of candidate regulatory interactions but rarely provide explicit confidence scores or credibility measures for these predictions. Moreover, existing methods lack mechanisms to incorporate biomedical literature for prioritizing TF–target relationships for experimental validation. In practice, researchers must manually search for evidence supporting each predicted TF–target relationship in the relevant cellular context. This process is increasingly impractical given the scale and growth of the literature (over 35 million PubMed abstracts and ~1 million new entries per year) and the largely unstructured reporting of findings (Alamro *et al*. 2024). As a result, the absence of automated, post-hoc evidence assessment remains a major barrier to translating inferred GRNs into testable hypotheses.

Automated text-mining efforts have extracted thousands of TF– target gene interactions from publications. For example, the ExTRI project identified over 53,000 TF–target relationships from PubMed abstracts (Vazquez *et al*. 2022). However, such resources function primarily as standalone text-mining databases and are not integrated into GRN inference pipelines as an automated post-hoc assessment step. As a result, researchers continue to struggle to determine which computationally inferred TF–target relationships are supported by existing evidence and should be prioritized for experimental validation through knockout or inhibition studies. This gap underscores a clear unmet need for automated frameworks that bridge GRN inference outputs with literature-based evidence assessment.

We introduce RAGulate, a novel framework that addresses this challenge by enabling post-hoc, literature-grounded verification of GRN predictions through a retrieval-augmented generation (RAG) strategy. In this approach, an LLM is “augmented” with a literature retrieval step through an external domain-specific knowledge database to ground its outputs in published evidence. Specifically, given a candidate regulatory edge or triplet (a TF–target gene pair provided some cell type/cell line) from a predicted GRN, RAGulate automatically retrieves relevant documents (e.g., PubMed abstracts or full-text papers) and prompts the LLM to generate an evidence-backed assessment of that regulatory relationship. Building on this, the system assigns a literature-grounded confidence score to each predicted relationship, reflecting the strength of supporting evidence. By design, RAGulate aims at mitigating the hallucinations often seen with unguided LLMs, reducing the risk of false associations (Nishisako *et al*. 2025).

To the best of our knowledge, no prior method exists for the assessment of GRN predictions via literature evidence in the post-hoc manner, making RAGulate a first-of-its-kind framework. It differs substantially from related RAGs in the biological research field. For example, GeneRAG, is a recently developed retrieval-augmented system that enhances LLM performance on gene-centric queries (Lin *et al*. 2024). GeneRAG demonstrated improved factual accuracy in tasks such as answering gene-related questions, cell-type annotation, and even gene interaction queries by leveraging external knowledge bases. However, GeneRAG is geared towards general Q&A and annotation tasks rather than systematically validating the output of GRN inference algorithms. Another recent approach, LLM4GRN, explored the use of prompting techniques (chain-of-thought reasoning and context injection) to guide GPT-4 in discovering gene regulatory networks (Afonja *et al*. 2025). While LLM4GRN demonstrated that an LLM can hypothesize plausible GRN connections (evaluated via synthetic data generation), it did not incorporate a retrieval step to ground those hypotheses in literature evidence. In fact, LLM4GRN relies on the model’s internal knowledge and provided context, which makes it susceptible to both knowledge gaps and hallucinations.

On the other hand, most applications of LLMs in genomics and bioinformatics have focused on addressing other important challenges. For example, automating cell-type annotation by using marker gene descriptions, or improving text-mining tasks with domain-specific foundation models like BioBERT (Lee *et al*. 2020) and PubMedBERT (Gu *et al*. 2021). These domain-trained Transformers have advanced biomedical named entity recognition and relation extraction (Alamro *et al*. 2024), but they are discriminative models and are not designed to generate natural-language verification for GRN predictions. In short, prior work has not tackled the specific challenge that RAGulate addresses: using an LLM with dynamic literature retrieval to provide post-hoc assessment of computationally inferred gene regulatory relationships. RAGulate thus fills a unique niche at the intersection of GRN inference and literature-based model explainability.

The significance of this work lies in addressing the crucial issue of LLM hallucinations and evidence retrieval in the context of GRNs, thereby increasing confidence in computational predictions. Key technical contributions include: (A) utilization of an external knowledge database CollectRI (cite here) of TF-target gene relationship that enables context-aware retrieval and evidence conditioning for regulatory assessment; (B) alias normalization of TF and gene symbols to enable robust, alias-aware query construction and retrieval, by incorporating the HUGO Gene Nomenclature Committee (HGNC) complete set of approved symbols and aliases (Tweedie *et al*. 2021); (C) the integration of a RAG pipeline that finds and leverages relevant publications on PubMed to assess each predicted regulatory interaction; (D) a method to quantify confidence in GRN connections in a context based on the presence (or absence) of supporting literature (including an algorithm for scoring interactions using retrieved evidence); and (E) an open-source, modular implementation that supports interchangeable retrieval backends and language models, facilitating deployment across datasets and compute environments.

Collectively, these contributions bridge the gap between in-silico network inference and real-world biological knowledge, ultimately helping to prioritize hypotheses and guide experimental validation efforts.

## 2 Methods

### 2.1 Overview of the RAGulate architecture

RAGulate is designed as a modular retrieval-augmented generation pipeline for post-hoc knowledge-based assessment of transcriptional regulation (Figures 1 & S1). At a high level, the system accepts the description of a transcription factor, a target gene, and a biological context as input and produces a confidence score for prioritizing TF–target regulation in the specified context, along with literature-grounded evidence. The pipeline comprises five key components: (1) a knowledge base of TF-target interactions annotated with context, compiled from the CollectRI database (Müller-Dott *et al*. 2023); (2) an alias-aware hybrid retrieval engine that first normalizes and expands TF and gene names using a curated alias map, then ranks PubMed articles relevant to the query using BM25 (Robertson *et al*. 1994), sentence-embedding; (3) a prompt assembler that injects the top retrieved passages and the TF–target–context triplet description into task-specific instruction templates (e.g., regulatory classification, evidence summarization); (4) an LLM that processes this instruction-plus-context prompt to produce a textual assessment and implicit confidence; and (5) a score combiner that integrates the language model’s outputs with retrieval-based support to produce a final probability of regulation. Each component is decoupled so that alternative retrieval models, alias maps, prompt templates or language models can be swapped in without affecting the rest of the system.

**Figure 1.**
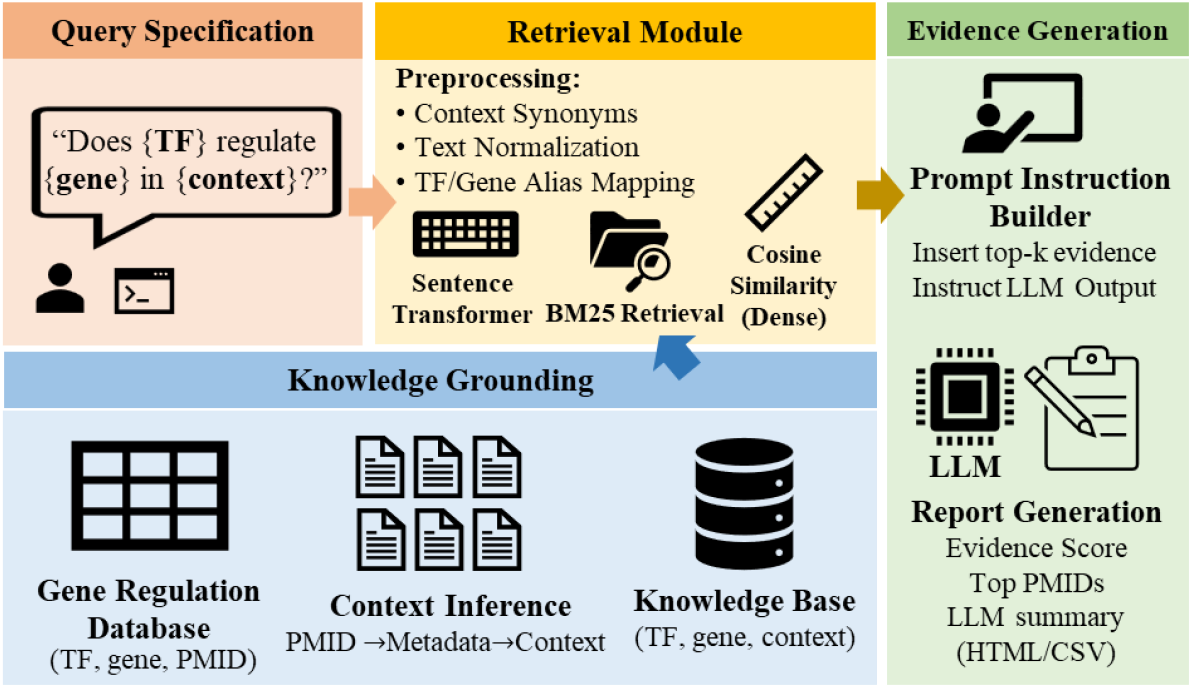
Overview of the RAGulate pipeline for context-aware transcriptional regulatory evidence retrieval and generation. RAGulate takes a structured query of the form “Does {TF} regulate {gene} in {context}?” and retrieves mechanistic evidence from PubMed for TF–gene–PMID triplets defined by the CollectRI network. The retrieval module preprocesses the query using context synonym expansion, text normalization, and TF/gene alias mapping before applying both dense (sentence-transformer cosine similarity) and sparse (BM25) retrieval. Retrieved abstracts are grounded with metadata-derived context labels to construct a TF–gene–context knowledge base. The top-k evidence is incorporated into a structured prompt, and an LLM generates a final report including evidence scores, supporting PMIDs, and a contextualized summary.

### 2.2. Data sources and knowledge base construction

#### 2.2.1. CollectRI regulatory interactions

RAGulate uses the CollectRI human regulatory database (Müller-Dott *et al*. 2023) as its primary source of TF-target interactions. Each row in the CollectRI table comprises a TF, its target gene, a signed weight indicating activation or repression and one or more PubMed identifiers (PMIDs) that support the interaction. We treat the weight as categorical to assign a regulation label: positive weights correspond to activation and negative weights correspond to repression. The set of unique PMIDs is extracted and cached on disk to avoid redundant network calls. PubMed titles, abstracts, and MeSH descriptors for each PMID are downloaded via NCBI Entrez using Biopython (Cock *et al*. 2009). Calls are rate-limited according to NCBI’s guidelines.

#### 2.2.2. Context inference and document construction

To enable context-aware retrieval, we infer biological context labels for each PubMed record using weak supervision based on MeSH descriptors and curated keyword lists. Evidence is mapped to a controlled vocabulary of eight canonical biological contexts, and each regulatory record is assigned one or more contexts with an associated confidence score. Records lacking sufficient signal are labeled as general. This procedure yields a structured, context-aware regulatory evidence corpus used for downstream retrieval and benchmarking (Supplementary Methods S1).

#### 2.2.3. Gold-standard benchmark

To evaluate retrieval performance and downstream binary regulatory classification (i.e., predicting whether a TF directly regulates a target gene in the queried context based on the evidence generated from the LLM), we construct a balanced gold-standard set of TF–target–context edges. Positive edges are sampled from CollecTRI by requiring at least one supporting PMID and a context score above the same threshold used during knowledge base construction. Negative edges are constructed by context-shuffling: for each context, we select TF–target pairs that are positive (i.e., supported by at least one PMID and passing the context-score threshold) in at least one other biological context but have no evidence in the current one, so that every negative example corresponds to a confirmed TF–target interaction that acts as a context-specific negative. This procedure produces negatives that are biologically plausible—true interactions but unsupported in the tested context— avoiding unrealistic random TF–gene pairings. We sample an equal number of positive and negative edges per context, capping the total at approximately 1,000 positives and 1,000 negatives. Each entry in the gold-standard file thus contains a TF, a target gene, a context label, a binary label (1 = evidence of direct regulation, 0 = no evidence) and a list of supporting PMIDs for positives, with an empty PMID list for negatives by construction.

### 2.3. Retrieval methods

For each TF–target–context triplet in the gold standard set, we construct a retrieval query of the form “{TF} regulates {gene} in {context}”. Here {TF} and {gene} denote symbol expressions that can include alias mapping, and {context} is either the canonical context label or its keyword-expanded variant. When alias-aware retrieval is enabled, we use the HGNC complete set of approved gene symbols (Tweedie *et al*. 2021) to map each TF and target symbol to a small alias set. These alias sets are incorporated into an OR-expanded query; for example, “(TP53 OR P53 OR LFS1) regulates (BAX OR BCL2) in immune”. We cap the number of alias terms (excluding the original symbol) to at most four per symbol.

We evaluate several retrieval strategies and measure their ability to recover the gold-standard PMIDs at different cut-offs (see tasks in Table 1).

**Table 1.** Summary of benchmarking tasks and evaluation settings in RAGulate.

| *Evaluation Stage* | **Benchmark Task** | **Description of Experiment** | **Configurations Compared** | **Metrics** |
| --- | --- | --- | --- | --- |
| *Retrieval (R)* | Effect of HGNC alias mapping on retrieval (Task 1) | Evaluate whether TF/gene synonym expansion improves lexical retrieval quality. | BM25<br>BM25 + HGNC alias mapping | Precision@k<br>Recall@k |
|  | Retriever comparison across lexical, dense, hybrid, and MMR variants (Task 2) | Compare retrieval effectiveness across all retriever families, including dense encoders and diversity-promoting re-rankers. | BM25<br>BM25 + aliases<br>BioBERT cosine<br>MiniLM cosine<br>BioBERT + MMR<br>MiniLM + MMR<br>Hybrid<br>Hybrid + aliases (RAGulate)<br>Vanilla RAG | Recall@k |
|  | Evidence-level PMID recovery for TF-target regulatory interactions (Task 3) | Quantify how well each pipeline retrieves correct supporting PubMed articles for gold-standard TF-target interactions. | BM25 + heuristic scoring<br>BM25 + LLM scoring<br>RAGulate flagship (Hybrid + aliases)<br>RAGulate BioBERT<br>RAGulate MiniLM<br>RAGulate MiniLM + MMR<br>Vanilla RAG | PMID precision<br>PMID recall<br>% correct PMID |
| *Augmentation (A)* | Impact of retrieval on LLM-based regulatory classification (Task 4) | Compare LLM classification performance with and without retrieval conditioning to assess the contribution of retrieved literature. | Direct LLM (no retrieval)<br>RAGulate (Hybrid retriever) for:<br>• BioGPT<br>• Mistral<br>• Llama-3.1<br>• Phi-3<br>• Qwen-2.5 | AUROC<br>AUPR |
| *End-to-end (RAG)* | End-to-end regulatory classification under alternative retrieval backends (Task 5) | Evaluate full RAG pipelines by varying only the retrieval backend to isolate its contribution to downstream classification accuracy. | BM25 + heuristic<br>BM25<br>RAGulate flagship (Hybrid + aliases)<br>RAGulate BioBERT<br>RAGulate MiniLM<br>Vanilla RAG encoder<br>RAGulate MiniLM + MMR | AUROC<br>AUPR<br>Accuracy |
| *Generation (G)* | Faithfulness and citation correctness of generated rationales (Task 6) | Assess whether model rationales remain grounded in retrieved evidence and whether cited PMIDs are correct. | RAGulate flagship<br>Vanilla RAG<br>LLM only | Faithfulness<br>Truthfulness<br>PMID precision<br>PMID recall |

#### 2.3.1. BM25 baseline

Our first baseline uses the Best Matching (Okapi BM25) implementation (Robertson *et al*. 1994) to rank all cached PubMed documents. Each document is tokenized and lower-cased; the query is similarly tokenized. BM25 estimates document relevance by combining term frequency, inverse document frequency, and document-length normalization. For a query *q* containing terms *t*, the BM25 score of a document *d* is:

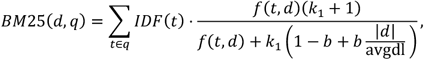

where *f*(*t, d*) is the term frequency of *t* in *d*, |*d*| is the document length, avgdl is the average document length in the corpus, *k*_1_ controls term-frequency saturation, and *b* controls length normalization.

We use default BM25 parameters (*k*_1_ = 1.5, *b* = 0.75) and return the top 50 documents (top_k) for each query. Retrieval performance is quantified by mean recall@k, precision@k, mean reciprocal rank (MRR) and mean average precision (MAP) averaged across all edges (see 2.5.1 for definitions).

#### 2.3.2. Vector-based retrieval in RAGulate

We encode titles and abstracts of all PubMed records using sentence embedding models. Two encoders are considered: (1) MiniLM (Reimers and Gurevych 2019; Wang *et al*. 2020), specifically, ‘all-MiniLM-L6-v2’ and (2) a BioBERT-derived sentence encoder fine-tuned for semantic textual similarity and natural language inference (Lee *et al*. 2020), specifically ‘pritamdeka/BioBERT-mnli-snli-scinli-scitail-mednli-stsb’. The query is similarly embedded, and cosine similarity between the query vector and every document vector is computed. The top 50 documents based on the similarity measure are returned. To improve diversity, we optionally apply maximum marginal relevance (MMR), re-ranking with λ = 0.5 or 0.7. MMR iteratively selects documents balancing relevance to the query against redundancy with already selected documents. For the query *q*, at iteration *i*, the next document *d*_*i*_ is chosen as:

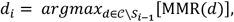

where:

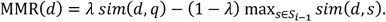

Here, 𝒞 is the candidate set, *S*_*i*−1_ is the set of already selected documents, and *sim*(·,·) is cosine similarity in embedding space. These retrieval configurations are denoted R1–R9 in our experiments (Task 2 in Table 1).

#### 2.3.3. Vanilla RAG and hybrid retrieval

As a reference we include the default LlamaIndex retriever (i.e., vanilla RAG) which uses its own embedding and Faiss index (Liu 2022). We also implement a hybrid retriever that combines BM25 and vector scores (R8). In this model we first obtain the top 300 BM25 candidates and the top 50 vector candidates, take the union, min–max normalize BM25 and vector scores, and rank the merged set by a weighted score (0.7 × BM25 + 0.3 × cosine similarity). The hybrid retriever optionally expands TF and target symbols with HGNC aliases; this configuration constitutes our “RAGulate flagship” model (R9) (Task 2 in Table 1). All retrieval runs are executed using identical top-k values.

### 2.4. Retrieval-augmented classification

#### 2.4.1. Prompt construction

For each edge in the gold-standard set we build a prompt to query an LLM. The prompt contains a question of the form: “Does TF X regulate target Y in context C?” followed by a context window comprising the top retrieved PubMed documents. Each document is rendered as “Title: … Abstract: … [PMID]”. We select at most 25 documents to respect the LLM’s input length limits and truncate abstracts to stay within a 4,096-token budget. Within each document, we include at most three extracted sentences (MAX_SENT_PER_PAPER = 3) to control redundancy and prompt length. Prompts instruct the model to answer “yes” or “no” and, where appropriate, include references to supporting PMIDs. For binary classification we cap generation to eight new tokens (GEN_MAX_NEW = 8); when a contextualized summary is requested, we allow up to 160 tokens (LLM_SUMMARY_MAX_NEW = 160). All prompts are generated deterministically based on retrieval results.

#### 2.4.2. LLM models and inference

We evaluate several off-the-shelf LLMs (Task 4 in Table 1): Bi-oGPT-Large (Luo *et al*. 2022), Mistral-7B-Instruct-v0.2 (Jiang *et al*. 2023), Meta-Llama-3-Instruct-8B (Grattafiori *et al*. 2024), Phi-3-mini-128k-instruct (Abdin *et al*. 2024) and Qwen-2.5-B-Instruct (Qwen *et al*. 2025). Models are accessed via the Hugging Face transformers library (Wolf *et al*. 2020) and run in half-precision on an NVIDIA DGX Spark GPU.

For binary regulatory classification, we use model-specific inference procedures aligned with each model’s training objective. For BioGPT-Large, we compute a token-level probability score from the model’s next-token distribution: given the assembled prompt, we extract the logits at the final input position, restrict the candidate set to the two tokenizer tokens corresponding to “yes” and “no” (including leading whitespace), and apply a softmax over these two logits. The resulting probability *P*(yes|prompt) is used directly as the continuous classification score, with a threshold defining the binary decision. In contrast, instruction-tuned LLMs (Mistral-7B-Instruct-v0.2, Meta-Llama-3-Instruct-8B, Phi-3-mini-128k-instruct, and Qwen-2.5-B-Instruct) are queried in deterministic generative mode, and the output is mapped to a binary label by parsing the first generated word (“yes” → 1, “no” → 0), with simple fallback heuristics when the response does not begin with an explicit yes/no token. This fallback heuristic procedure is keyword-based: if the first generated word is not an explicit “yes” or “no,” we assign a positive label when “yes” appears anywhere in the response and “no” does not, assign a negative label when “no” appears and “yes” does not, and otherwise assign an uncertain score of 0.5 (e.g., when both tokens appear or neither appears).

#### 2.4.3. Combining LLM and retrieval support

For RAGulate classification we combine the raw LLM probability (*L*) with a retrieval support score (S) derived from the ranks of retrieved documents. The combined confidence score uses a mixture of a linear term and a multiplicative term

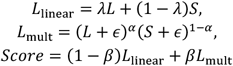

where *λ* = 0.5, *α* = 0.5, *β* = 0 and *ϵ* = 10^−8^ in our experiments. When *β* = 0, the multiplicative term is disabled, and the score reduces to a simple average of LLM and retrieval support. Edges are classified as positive when the score exceeds a threshold of 0.5.

### 2.5. Evaluation and statistical analysis

#### 2.5.1. Retrieval metrics

For retrieval-only experiments (R1–R9), we evaluate performance using standard information-retrieval metrics, including recall@k and precision@k for k ∈ {1, 3, 5, 10, 20, 50, 100}, mean reciprocal rank (MRR), and mean average precision (MAP), computed against gold-standard PMIDs for each TF–target–context query (Tasks 1–3 in Table 1). To assess the effect of HGNC alias expansion, confidence intervals for recall@k and precision@k are estimated using non-parametric bootstrapping.

Recall@k measures the fraction of gold-standard PMIDs retrieved within the top k results, while precision@k measures the proportion of retrieved documents that are relevant. For a single query instance (a TF–target–context triplet) with gold-standard PMID set *G* and retrieved list of PMIDs *R* we define:

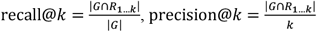

where *R*_1..*k*_ = {*R*_1_, …, *R*_*k*_} is the set of the top-*k* retrieved PMIDs.

MRR is the mean of the reciprocal ranks of the first relevant document, and MAP averages precision at each rank where a relevant document is found. The average precision (AP) for one edge is:

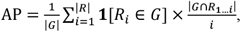

where **1**[*R*_*i*_ ∈ *G*] denotes the indicator function, equal to 1 when the retrieved item at rank *i* is relevant and 0 otherwise. The mean reciprocal rank is:

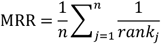

where rank_*j*_ is the rank of the first relevant document for the j-th query instance; *n* is the number of evaluation instances in the test set.

To compare retrieval with and without HGNC alias expansion, we use non-parametric bootstrapping (1,000 replicates) over TF–target–context query instances to compute 95% confidence intervals on recall@k and precision@k.

#### 2.5.2. Classification metrics

For regulatory classification, we evaluate predictive performance using AUROC, AUPR, accuracy, and maximum F1 score against gold-standard labels (Task 5 in Table 1). Thresholds are selected to maximize F1, and statistical significance between direct LLMs and their retrieval-augmented counterparts is assessed using paired, non-parametric permutation tests. Full metric definitions and testing procedures are described in Supplementary Methods S2.

#### 2.5.3. Evidence faithfulness and truthfulness

Beyond binary correctness, we evaluated whether LLM-generated rationales remain grounded in retrieved literature and whether cited PMIDs are correct with respect to the gold-standard evidence. Each evaluation instance corresponds to a single TF–target–context triplet or edge.

For each edge *e*, we extract the set of PMIDs mentioned in the generated summary, *P*_*pred*_(*e*), using a PMID-matching parser (e.g., patterns such as “PMID: #######” and bare numeric PubMed IDs when unambiguous). We also record (i) the set of PMIDs returned by the retriever, *P*_*ret*_ (*e*), and (ii) the gold-standard supporting PMID set, *P*_*gold*_(*e*).

We define a generated summary as *hallucinated* if it cites at least one PMID that was not present in the retrieved evidence set:

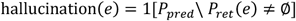

where 1[·] is the indicator function. The hallucination rate is the fraction of edges with hallucination(*e*) = 1. We report faithfulness across all evaluated edges *N* as:

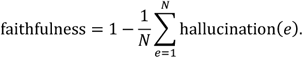

This metric penalizes any instance in which the model invents (hallucinates) a PMID not supported by retrieved context, thereby quantifying whether rationales remain constrained to the retrieval window.

We define an edge as *strictly correct* if the model cites at least one PMID and all cited PMIDs are contained in the gold standard evidence set:

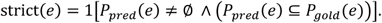

Above, 1[·] is the indicator function and ∧ denotes the logical AND operator. Furthermore, ∅ is an empty set, and we treat an empty predicted set *P*_*pred*_(*e*) = ∅ as providing no evidence. We report truthfulness as the mean strict correct rate across all the same *N* evaluated edges:

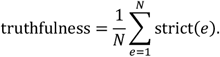

This definition is intentionally conservative: an output that cites no PMIDs is not counted as truthful, and any spurious PMID (even if some are correct) fails strict correctness.

To provide graded evidence overlap, we also compute PMID-level precision and recall per edge:

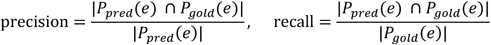

with the convention that precision(e) = 0 when *P*_*pred*_(*e*) = ∅. We summarize these quantities by their mean across edges.

#### 2.5.4. Implementation details

All code is written in Python 3.10 and structured into modular components for data loading, retrieval, LLM prompting, evaluation, and plotting. PubMed metadata and sentence embeddings are cached locally to ensure reproducibility. Experiments are seeded for reproducibility (seed = 42) and run on an NVIDIA DGX Spark workstation. The notebooks used to construct the knowledge base and run the benchmark are available at https://github.com/YDaiLab/RAGulate.

## 3 Results

We evaluated RAGulate using a curated benchmark from CollectRI, constructed from thousands of interactions (Figure 2A), where each TF– target pair was paired with context-inferred evidence (Figure 2B-C) and context-shuffled negatives. We first assessed retrieval performance, followed by downstream classification and evidence faithfulness analyses (Table 1). Results are summarized below.

**Figure 1a.**
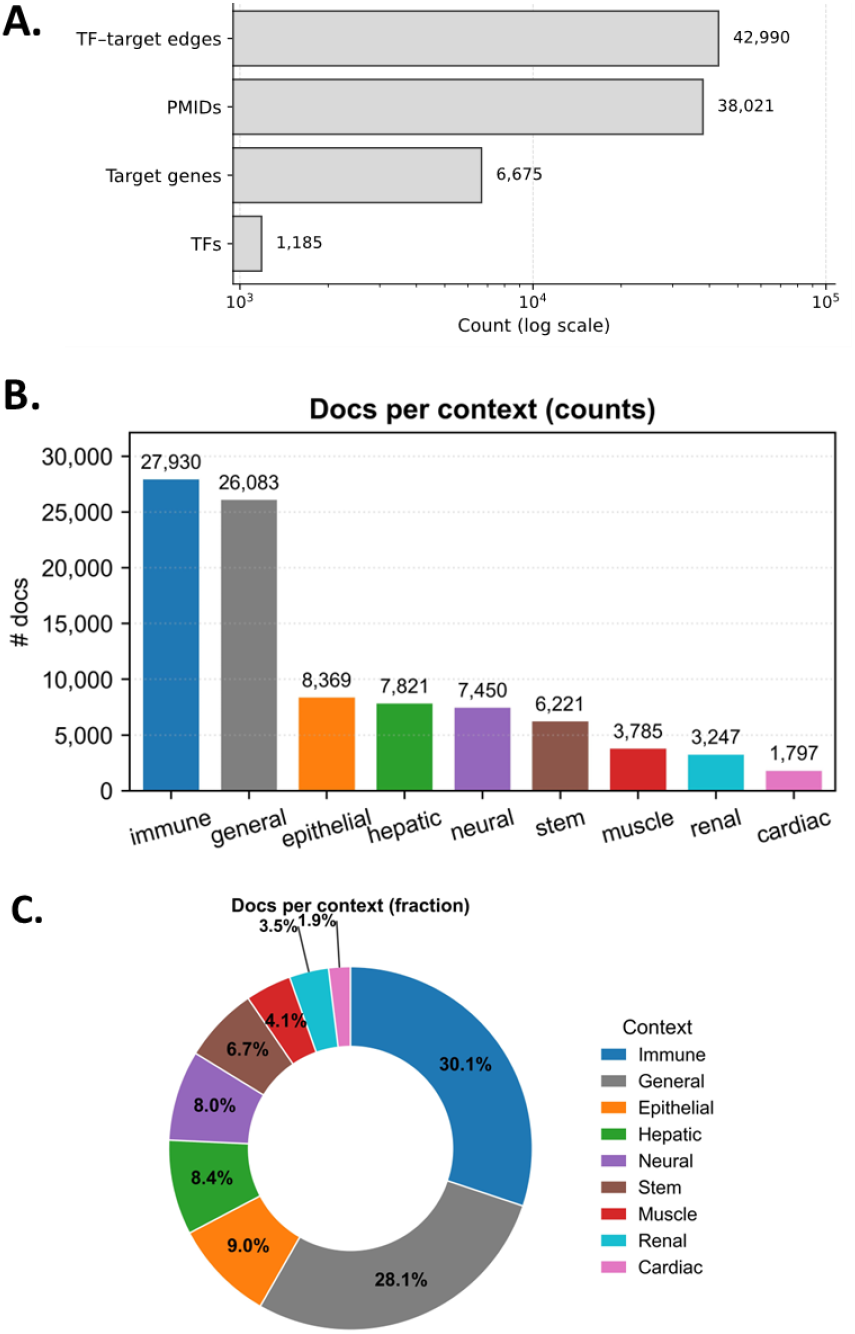
Distribution of inferred biological contexts across CollectRI regulatory evidence. **(A)** CollecTRI (human) dataset size. **(B)** Bar plot showing the number of PubMed records assigned to each biological context after applying the metadata-based context inference module to CollectRI TF-gene-PMID entries. Immune and general contexts dominate the evidence landscape, followed by epithelial, hepatic, and neural contexts, with smaller contributions from stem, muscle, renal, and cardiac literature. **(B)** Donut plot showing the proportional representation of each biological context. Immune and general contexts account for the majority of evidence, while epithelial, hepatic, neural, stem, muscle, renal, and cardiac contexts appear with progressively lower fraction.

### 3.1 Context inference organizes CollectRI evidence into biologically meaningful categories

We first applied the metadata-based context inference module to all CollectRI TF–gene–PMID records. Each publication was assigned to one of eight canonical biological contexts using MeSH voting and curated keyword lists. This procedure generated a structured regulatory evidence landscape, where immune and general literature constituted the largest fractions, followed by epithelial, hepatic, neural, and stem contexts, with smaller contributions from muscle, renal, and cardiac categories (Figure 2B–C). These distributions highlight the uneven availability of regulatory evidence across biological domains and motivate the need for retrieval methods capable of adapting to heterogeneous context densities.

### 3.2 Alias normalization substantially improves lexical retrieval performance

We next evaluated the effect of HGNC alias expansion on BM25 retrieval (Table 1, Task 1). Alias-aware queries consistently increased both precision@K and recall@K relative to raw symbol queries, particularly at small cut-offs where symbol ambiguity is most detrimental to ranking quality (Figure 3A–B). Recall improvements approached the sub-string-matching upper bound, demonstrating that alias expansion resolves a major proportion of missed matches arising from heterogeneous gene and TF nomenclature. These findings establish symbol normalization as a critical preprocessing step for literature-based regulatory inference.

**Figure 1b.**
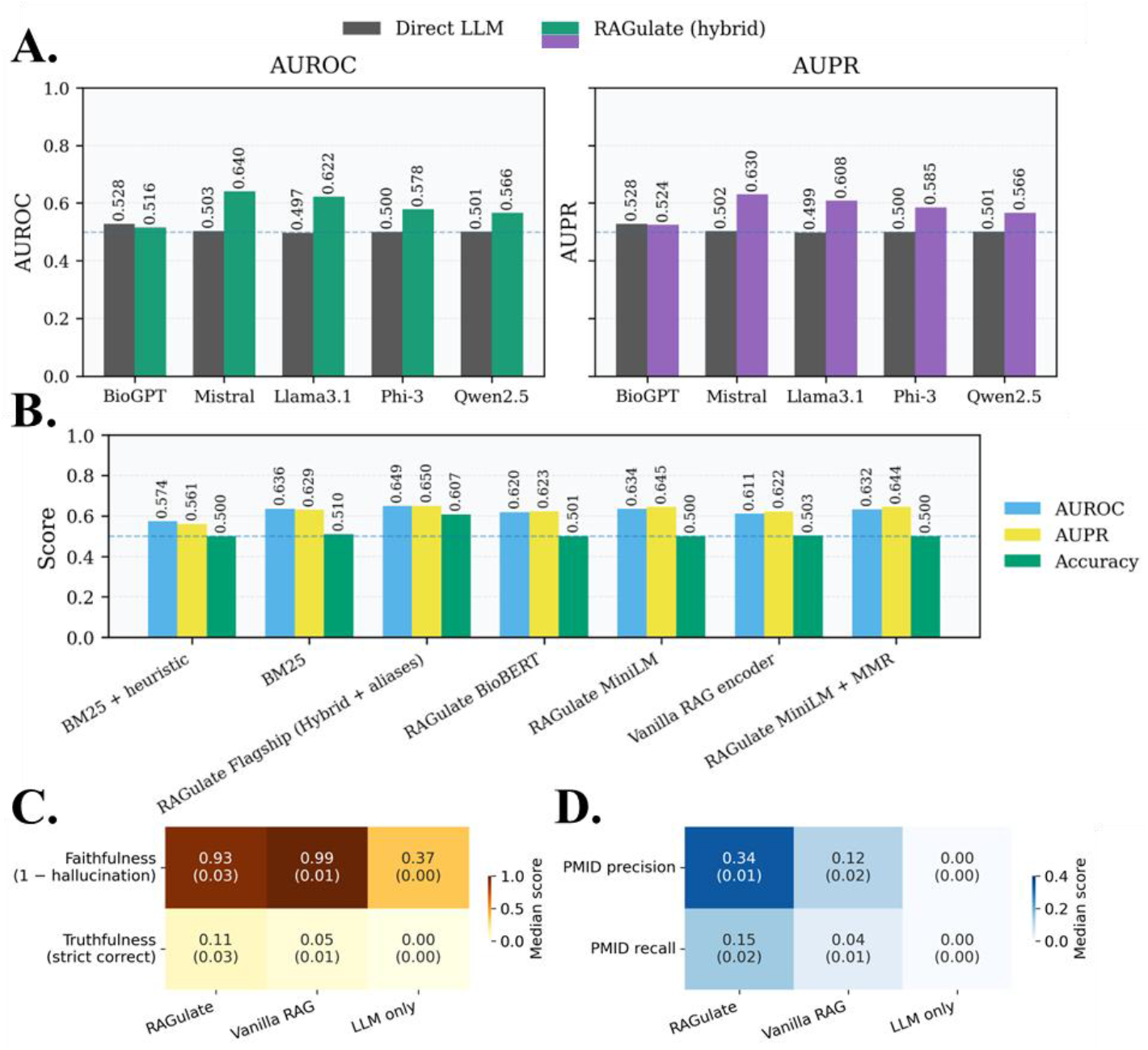
Benchmarking generation and end-to-end evidence integration in RAGulate. **(A)** AUROC and AUPR for direct LLMs versus RA-Gulate (hybrid retrieval + generation) across five foundation models, showing consistent performance gains when grounded with retrieved evidence. **(B)** End-to-end classification performance (AUROC, AUPR, Accuracy) across retrieval pipelines, demonstrating that hybrid and alias-augmented RAGulate configurations achieve the strongest predictive performance. **(C)** Faithfulness (1 − hallucination) and strict truthfulness of generated summaries across pipelines. RAGulate substantially reduces hallucination rates compared to vanilla RAG and direct LLM prompting. **(D)** PMID-level evidence benchmarking, measuring median precision and recall of cited evidence. RAGulate achieves the highest fidelity in recovering correct PMIDs among the retrieved and generated outputs.

### 3.3 Hybrid retrieval outperforms sparse and dense pipelines across all recall thresholds

To assess retrieval beyond lexical matching, we compared BM25, BioBERT and MiniLM dense encoders, maximum-marginal-relevance (MMR) variants, vanilla RAG retrieval, and the hybrid BM25+dense retriever used in RAGulate (Table 1, Task 2). Recall@20 comparisons (Figure 3C) and full recall@K curves (Figure 3D) show that dense retrieval improves evidence recovery over sparse methods but often retrieves redundant documents. MMR re-ranking mitigates redundancy and improves mid-range recall. Across all configurations, hybrid retrieval—which merges BM25 and dense candidates—achieved the highest recall, and alias expansion further amplified this advantage. These results identify hybrid retrieval with alias normalization as the most effective strategy for retrieving contextually relevant biomedical evidence.

### 3.4 Hybrid pipelines improve evidence-level PMID recovery for TF–target interactions

We then evaluated the ability of each retrieval configuration to recover the correct supporting PMIDs associated with gold-standard TF– target interactions. These configurations differ in retrieval signal (lexical, dense, or hybrid), encoder choice, and scoring strategy (Table 1, Task 3).

RAGulate flagship (Hybrid+Alias retriever and MiniLM encoder) achieved the highest rates of exact-match and top-k recovery, outperforming purely lexical, purely dense, and vanilla RAG pipelines (Figure 3E). This confirms that hybrid retrieval, augmented with alias mapping, is particularly effective at identifying the specific studies that underlie curated regulatory relationships.

### 3.5 Retrieval conditioning improves regulatory classification across diverse LLM architectures

To determine whether retrieved evidence improves model-based classification, we compared five LLMs: BioGPT (Luo *et al*. 2022), Mistral-7B (Jiang *et al*. 2023), Llama-3.1-8B (Grattafiori *et al*. 2024), Phi-3 (Abdin *et al*. 2024), and Qwen-2.5 (Qwen *et al*. 2025), when prompted directly versus when conditioned on the top retrieved documents from RAGulate (Table 1, Task 4). In every case, retrieval conditioning increased AUROC and AUPR (Figure 4A), indicating that LLMs benefit from explicit exposure to relevant literature even when they possess substantial biomedical domain knowledge. These improvements demonstrate the importance of grounding LLM predictions in retrieved evidence rather than relying solely on parametric memory.

**Figure 2.**
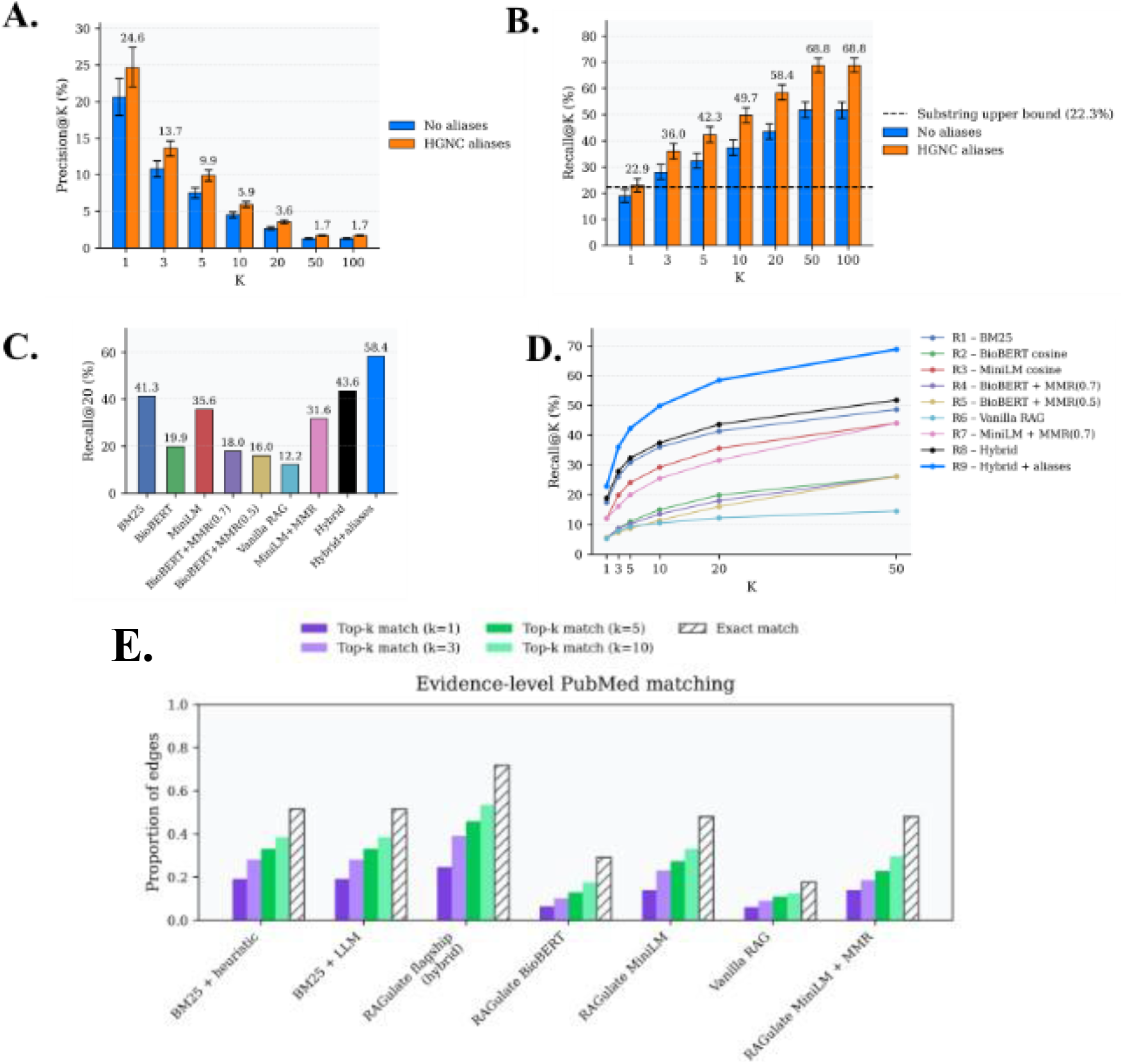
Benchmarking retrieval and evidence-matching components of RAGulate. **(A)** Precision@K with and without HGNC alias mapping, showing improved precision across all cutoffs when aliases are incorporated. **(B)** Recall@K with and without HGNC alias mapping; alias expansion substantially increases recall and approaches the substring-matching upper bound at large K. **(C)** Recall@20 across retrieval configurations (sparse, dense, hybrid, and hybrid + aliases), demonstrating superior evidence recovery for hybrid pipelines. **(D)** Recall@K curves for nine retrieval pipelines, comparing sparse, dense, hybrid, and hybrid-MMR variants; hybrid + alias mapping achieves the highest retrieval performance. **(E)** Evidence-level PubMed matching for TF–target interactions across baseline and RAGulate pipelines, showing improved exact-match and top-k recovery for hybrid and alias-augmented configurations.

### 3.6 End-to-end pipelines demonstrate that retrieval quality governs predictive accuracy

We further benchmarked full RAG pipelines in which only the retrieval module differed (Table 1, Task 5). Pipelines using hybrid retrieval produced the strongest predictive performance across AUROC, AUPR, and accuracy (Figure 4B). Dense-only, sparse-only, and default RAG retrieval consistently underperformed relative to hybrid configurations. This establishes retrieval quality emerged as the dominant factor governing classification accuracy, outweighing differences across LLM architectures.

### 3.7 RAGulate reduces hallucinations and improves evidence faithfulness in generated rationales

We quantified the grounding of generated explanations by measuring hallucination rate (faithfulness) and strict correctness of cited evidence (Table 1, Task 6). RAGulate produced the most faithful summaries, showing markedly reduced hallucination compared to vanilla RAG and direct prompting (Figure 4C). PMID-level evaluation similarly showed that RAGulate achieved the highest median precision and recall of cited evidence (Figure 4D). These results indicate that accurate retrieval not only improves classification but also constrains the LLM to generate explanations that accurately reflect the supporting literature.

### 3.8 Case study: RAGulate yields interpretable, evidence-linked regulatory assessments

Finally, we illustrate RAGulate’s utility on real TF–target–context queries (Figure S2). Specifically, we show the queries (TBX21, GZMB, NK cell) and (PAX5, CD19, naïve B cell). For each query, the system retrieves the most relevant PubMed passages, highlights evidence supporting the regulatory relationship, and generates an LLM summary linking TF function, target gene behavior, and context-specific biology. The resulting table shows both high-scoring interactions, along with the precise PMIDs and rationale underlying the final decision. These examples demonstrate how RAGulate produces transparent, literature-grounded regulatory assessments that can be directly inspected by researchers.

## 4 Discussion

This study introduces RAGulate, a modular retrieval-augmented generation framework for post-hoc assessment of transcription factor–target regulation using the biomedical literature. Although numerous resources curate regulatory interactions, their evidence is heterogeneous, context-dependent, and scattered across thousands of publications. By coupling alias-aware retrieval, dense and sparse ranking, and LLM-based reasoning over an external knowledge base of PubMed abstracts, RAGulate provides a unified pipeline that can recover relevant literature, score regulatory plausibility, and generate transparent explanations.

A central observation from our evaluation is that traditional lexical retrieval alone is insufficient for recovering the breadth of evidence needed for regulatory assessment. Gene and TF symbols are often ambiguous, inconsistently reported, or absent from abstracts, which degrades BM25 performance. Alias expansion mitigates this issue and yields substantial improvements in recall and precision, particularly at small k, where mis-tokenization has the greatest impact. These gains approached the substring-matching upper bound, underscoring the importance of systematic symbol normalization for literature-based regulatory inference.

Dense retrieval further improved evidence recovery by capturing semantic similarity that is not detectable through lexical matching. However, dense encoders also tend to retrieve near-duplicate documents and exhibit limited robustness to alias variation. Our results show that maximum-mar-ginal-relevance re-ranking partially addresses redundancy but does not eliminate it. In contrast, hybrid retrieval that merges sparse lexical and dense semantic candidates consistently outperformed all individual strategies across recall thresholds. These findings indicate that hybridization is necessary to balance BM25’s precision with dense models’ ability to capture conceptual relevance, and suggest that future biomedical retrieval systems should default to hybrid architectures.

Retrieval quality directly influenced downstream regulatory classification. Across five LLMs, conditioning on retrieved evidence improved AUROC and AUPR relative to direct classification, demonstrating that even domain-adapted models such as BioGPT benefit from explicit exposure to relevant literature. This effect was consistent across model sizes and architectures, suggesting that retrieval plays a more critical role than parametric memory for this task. End-to-end benchmarks further confirmed that retrieval performance (not the LLM) is the primary determinant of classification accuracy. Pipelines using hybrid retrieval achieved the strongest results, whereas sparse-only, dense-only, and default RAG retrieval lagged behind.

Beyond binary correctness, RAGulate substantially improved evidence faithfulness. The system produced fewer hallucinated statements and achieved higher PMID-level precision and recall compared with direct prompts and vanilla RAG. This was especially notable given the importance of citation correctness for scientific workflows. Case studies illustrated how RAGulate retrieves specific mechanistic evidence, highlights supporting text spans, and generates concise contextual summaries that link TF function, target gene behavior, and biological setting. These interpretable outputs offer a transparent alternative to black-box predictions and may serve as literature-aided annotations for regulatory databases.

Several limitations merit consideration. First, context inference relies on MeSH terms and curated keyword lists, which may not capture fine-grained or non-canonical biological conditions. Expanding the context vocabulary or incorporating supervised context classifiers may improve specificity. Second, our benchmark focuses on interactions present in CollectRI; interactions absent from the knowledge base cannot be evaluated for evidence recovery, and LLM classification on such edges may require a more conservative interpretation. Third, the combination of LLM and retrieval scores uses fixed mixture weights; adaptive or learned weighting strategies could further optimize classification. Finally, although retrieval conditioning improves performance, LLM outputs still depend on the model’s internal reasoning, and misinterpretations of evidence remain possible. Future extensions could incorporate citation-verification objectives or chain-of-thought supervision to strengthen grounding.

Overall, the results demonstrate that retrieval is a key determinant of reliable literature-grounded inference. By integrating retrieval and generation within a modular framework, RAGulate provides a flexible foundation for evidence-based interpretation of regulatory interactions and can be extended to other modalities of biological knowledge extraction.

## 5 Conclusion

RAGulate provides a systematic approach for validating transcriptional regulation by combining alias-aware retrieval, hybrid ranking, and LLM-based reasoning. The architecture consistently improves evidence recovery, enhances regulatory classification across diverse LLMs, and reduces hallucination in generated rationales. Hybrid retrieval with alias expansion emerged as the most effective strategy, revealing the central role of retrieval quality in literature-grounded inference. Through its modular design and interpretable outputs, RAGulate offers a practical tool for researchers seeking to verify regulatory hypotheses, curate database entries, or explore context-specific regulatory mechanisms directly from biomedical literature.

## Supplementary data

Supplementary data are available at *Bioinformatics* online.

## Conflict of interest

None declared.

## Funding

This work was supported by NVIDIA Corporation through its academic grant program via the provision of an NVIDIA DGX Spark system.

## Data availability

The latest HGNC alias list was downloaded from https://storage.googleapis.com/public-download-files/hgnc/tsv/tsv/hgnc_complete_set.txt.

The data underlying this article are available on GitHub at https://github.com/YDaiLab/RAGulate, and archived at https://doi.org/10.5281/zenodo.18320507.

## Notes

https://github.com/YDaiLab/RAGulate

### Competing Interest Statement

The authors have declared no competing interest.

