## Supplementary Material for "RAGulate: Retrieval-Augmented Generation for Post-hoc Literature-Grounded Regulatory Assessment"

Contents:


**Supplementary Methods S1**: Metadata-based biological context inference

**Supplementary Methods S2**: Classification metrics

**Supplementary Figure S1**: Component-level architecture of a retrieval-augmented generation (RAG) framework

**Supplementary Figure S2**: Case study demonstrating RAGulate's end-to-end regulatory assessment

**Supplementary Methods S1: Metadata-based biological context inference**

We define eight canonical biological contexts (immune, neural, cardiac, hepatic, renal, muscle, epithelial and stem). Each MeSH descriptor associated with a PubMed record is mapped to one or more contexts using a manually curated dictionary. To capture contextual information when MeSH annotations are absent or ambiguous, we supplement this mapping with per-context keyword lists.

For each record, we compute a context score by summing MeSH votes (weight = 1.0) and keyword votes (weight = 0.5) for each context and normalizing the resulting scores to the range [0,1]. We retain up to two contexts with scores ≥ 0.3. Records that do not exceed this threshold for any context are assigned to a "general" category.

For each CollectRI TF–target–PMID triplet, we generate a concise sentence describing the interaction and attach metadata including the TF, target gene, regulatory sign, assigned context label, per-context confidence score, and the full context score vector prior to thresholding.

**Supplementary Methods S2: Classification metrics**

For classification experiments we treat the gold-standard label as the true class and the combined score or LLM score as the prediction. We report area under the receiver-operating characteristic curve (AUROC), area under the precision–recall curve (AUPRC), the maximum F1 score and the corresponding probability threshold and accuracy. Given true positives (TP), false positives (FP), true negatives (TN) and false negatives (FN) at a chosen threshold, precision P and recall R are:

$$\mathrm{P} = \frac{\mathrm{TP}}{\mathrm{TP+FP}} \text{ and } \mathrm{R} = \frac{\mathrm{TP}}{\mathrm{TP+FN}}.$$

The F1 score is their harmonic mean:

$$F1 = 2\frac{P \times R}{P + R},$$

and accuracy is (TP + TN)/(TP + FP + TN + FN). We choose the threshold that maximizes F1 over the benchmark instances. When comparing direct LLMs against their RAGulate counterparts we use paired permutation tests (100 permutations) to assess whether performance differences are significant.

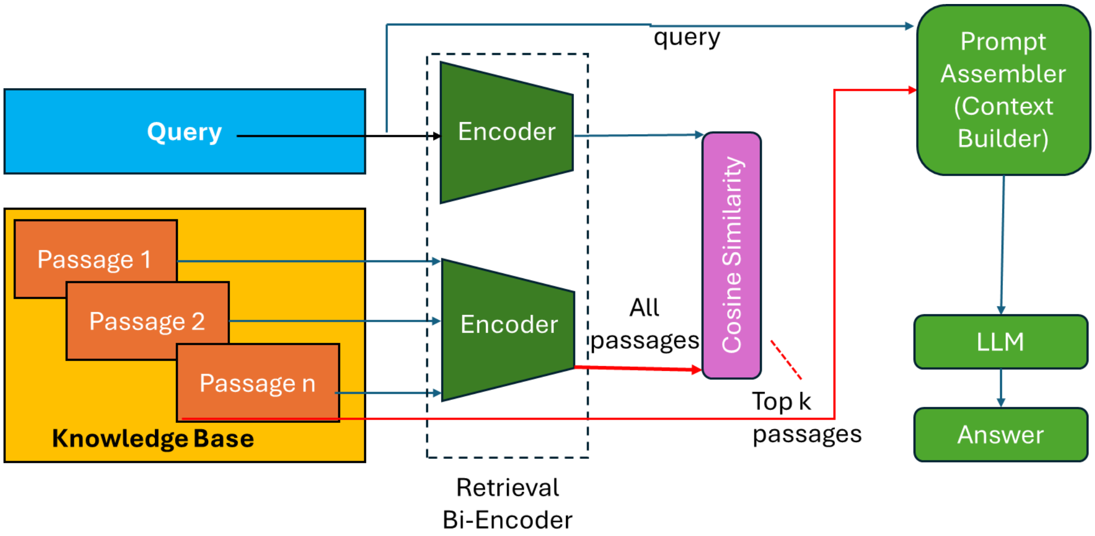


**Figure S1. Component-level architecture of a retrieval-augmented generation (RAG) framework.** The system consists of independent encoders for queries and knowledge-base passages, cosine similarity–based retrieval, a prompt assembly module, and an LLM that produces the final response.

| tf | target | context | score | evidence_highlighted | llm_summary |
|---|---|---|---|---|---|
| TBX21 | GZMB | NK cell | 0.864883 | [PMID 25352127] We subsequently found that TOX2 was independent of ETS-1 but could directly upregulate the transcription of **TBX21** (encoding T-BET). [PMID 25352127] Given the essential function of T-BET in **NK cell** differentiation, TOX2 therefore plays a crucial role in controlling normal **NK cell** development by acting upstream of **TBX21** transcriptional regulation. [PMID 24752800] Moreover, during the differentiation process NK cells gradually display increasing expression of IFNG and **TBX21** (encoding T-bet) transcripts and demethylation at the IFNG promoter. … (6 more sentences) Show more | Transcription factor TBX21, also known as T-bet, is essential for NK cell differentiation and is upregulated during the process. The upregulation of TBX21 is influenced by TOX2, which directly activates TBX21 transcription (PMID 25352127). However, the regulation of TBX21 remains incompletely defined, and other factors such as Notch signaling and Yin-Yang 1 binding to the TBX21 promoter may also play a role (PMIDs 15991363, 21272048). The evidence is a mix of direct and indirect. |
| PAX5 | CD19 | naive B cell | 0.865013 | [PMID 8639790] Recent phenotypic analysis of plasma cells showed that normal plasma cells do express the B-cell lineage-specific molecule **CD19**, but their malignant counterpart (myeloma cells) are **CD19**-. [PMID 8639790] To clarify the meaning of loss of **CD19** antigen on myeloma cells, we first compared the expression of **CD19** and Pax-5 genes among B cells, normal plasma cells, myeloma cell lines, and primary myeloma cells, because the Pax-5 gene was reported to encode the transcriptional factor, B-cell-specific activating protein (BSAP), necessary for **CD19** gene expression. [PMID 8639790] Neither **CD19** nor Pax-5 mRNA could be detected in those primary myeloma cells and cell lines, whereas normal plasma cells did express both **CD19** and Pax-5 mRNA. … (7 more sentences) Show more | PAX5, the transcription factor that encodes the B-cell-specific activating protein (BSAP), is crucial for the expression of the B-cell lineage-specific molecule CD19 in naive B cells. However, the correlation between the level of PAX5/BSAP expression and that of CD19 is not clear-cut, as some studies have failed to find a direct correlation between the two. [PMID 9545244] is the most direct evidence, showing that CD19 synthesis is critically dependent on the BSAP protein concentration. |

**Figure S2. Case study demonstrating RAGulate's end-to-end regulatory assessment.** Example TF–target–context queries processed by RAGulate, showing retrieval-highlighted PubMed evidence, final regulatory scores, and LLM-generated summaries. Retrieved sentences linking the TF and target are highlighted, and supporting PMIDs are shown explicitly. These examples illustrate how RAGulate integrates hybrid retrieval and generative modeling to produce interpretable, literature-grounded regulatory predictions.
